# Nutrient content estimation of the world’s fishes

**DOI:** 10.64898/2026.05.19.726181

**Authors:** M Aaron MacNeil, Eva Maire, James PW Robinson, Nicholas AJ Graham, Philippa J Cohen, Maria LD Palomares, Christina C Hicks

**Author notes:** Corresponding author(s): Aaron MacNeil.

## Abstract

Seafood nutrients from global fisheries are of increasing importance for research and policy in food security and nutrition. As the chemical composition of fish is determined by what they eat, their energetic demands, and the environment in which they live, nutrient content reflects aspects of physiology and life history, ecological and environmental traits, as well as evolutionary history. Here we present data from Bayesian model estimates of 12 key nutrients (calcium, iron, phosphorus, magnesium, selenium, zinc, vitamin A, vitamin B9, vitamin B12, vitamin D, omega-3 fatty acids, and protein) in wild fish, using a database of reported nutrient content for freshwater and marine species. We then predict the nutrient content of 5588 fish species with traits available from FishBase. We compare our previous model using traits alone with a new model of both traits and phylogeny, and present the data, code, and predictions for models coded in PyMC. These models and predictions, made freely available through FishBase, can be used to explore the historical, current, and future nutrient content of fisheries catch.

## Background & Summary

Micronutrient deficiencies are responsible for approximately one million premature deaths each year, along with a diversity of physical and cognitive developmental problems, making sufficient supply of nutritious foods a critical component of global food and nutrition security (Haddad et al. 2016). Fish have been identified as being a rich source of bioavailable macro- and micronutrients and are increasingly recommended for healthy diets (Golden et al. 2021). Recent analyses have demonstrated that strategies to alter production and distribution of aquatic foods caught in marine and inland fisheries have the potential to help address food insecurity for millions of people (Rittenschober et al. 2013, 2016, Hicks et al. 2019). However, standardized nutritional information for most species remains difficult to access. Analyses of nutrient content typically publish data for specific nutrients (*e*.*g*., protein, zinc) and focus on one or a few species per study, using different sampling and analytical methods. Nutrient content information for the thousands of fish caught and eaten globally have remained unavailable, limiting scientific exploration of the distribution of nutrient concentrations, yields from fisheries, and the development of nutrition-based fisheries policies.

Published nutrient studies do, however, represent diverse species that are targeted across the world’s fisheries, and contain data on multiple nutrients that are essential in diets. Statistical analysis of variation in these nutrient datasets can provide standardized estimates of fish nutritional content, for species with existing sample data as well as data-deficient species (FAO 2017). Theory suggests that variation in fish nutrient content should be reflected in three interrelated sets of attributes relating to their physical traits, environment, and evolutionary history. Fish gain nutrients from their diet (Galbraith et al. 2019) and so nutrient content will be influenced by their trophic position and feeding ecology. Fish morphology and physiology is also strongly related to diet, with attributes such as mouth structure, body shape, metabolism, and body size reflecting where and what they eat. In addition, nutrients and minerals available within a given food web are affected by environmental variation and the characteristics of primary producers that differ by depth, thermal range, and marine realm (Behrenfeld et al. 2006, Tagliabue et al. 2017). Lastly, the position of fish species within their phylogenetic lineage implies stronger correlation in nutrient content among more closely related species (Vaitla et al. 2018).

Conventional wisdom would suggest that correlations among related species in terms of both nutrient content and traits would imply that models developed to predict nutrient content should use either phylogeny (Vaitla et al. 2018) or traits (Hicks et al. 2019) as covariates, due to potential multicollinearity. Yet multicollinearity is only a problem for interpretation of model coefficients, not prediction (McElreath 2020), and affects some estimation methods more than others. Gradient-based algorithms of the kind used for Bayesian modelling have little difficulty with even highly correlated parameters, meaning that the most accurate predictions would likely come from models including both phylogeny and traits. Here we develop statistical models for estimating nutrient content of fishes globally, incorporating covariate information from phylogeny and traits, as well as potential sampling effects among studies. We evaluate model performance relative to our previously published model (Hicks et al. 2019) and use the most accurate models out-of-sample to produce estimated content of calcium, iron, phosphorus, magnesium, selenium, zinc, vitamin A, vitamin B12, omega-3 fatty acids, and protein for 5588 of the world’s fishes. The observed and estimated nutrient estimates are made freely available through the online database FishBase (Froese and Pauly 2010).

## Methods

### Data collection

Nutrient data was collected from three key sources into a database of 5,759 measurements of various nutrient compositions from 731 species. Data sources included: (1) a Thompson Reuters Web of Science search for published fish nutrient values between 1979 and 2022, using the search terms ‘content’ or ‘compos*’, and ‘nutrition* NEAR content NEAR fish* AND Marine*’; (2) the FAO/INFOODS food composition for biodiversity databases^3–5^ produced by the Food and Agriculture Organization (FAO) of the United Nations; and (3) additional key informant sources of robust grey literature, developed through snowball questioning of nutrition researchers. We retained sources that were in English, were fully traceable, from wild-caught fish, where scientific names were provided, where quantitative analytical data were reported in unequivocal matrix and units, analyses conducted on fresh samples, reported as whole body, fillet, muscle, or ‘edible portion’. Initial data collection took place in 2016, focused on marine finfish, this was updated in 2020 and 2022, and broadened to include freshwater species.

From these sources we extracted quantitative information on 12 key nutrients considered essential to human health, 6 minerals: calcium, iron, magnesium, phosphorus, selenium, zinc, 4 vitamins: vitamin A, vitamin B9, vitamin B12, vitamin D, and two macronutrients: protein, and for fats we separately recoded: poly unsaturated fatty acids (PUFA), total omega-3 fatty acids, total omega-6 fatty acids, eicosapentaenic acid (EPA) and docosahesaenoic acid (DHA) . We used total omega-3 fatty acids for all subsequent fatty acid analyses. Values were standardized to μg, mg, or g per 100g, depending on the unit relevant to dietary intake, and differences in sample characteristics (wet, dry, whole etc.) were recorded for subsequent modelling of potential bias. Once compiled, we had data for 12 nutrients (calcium, iron, phosphorus, magnesium, selenium, zinc, vitamin A, vitamin B9, vitamin B12, vitamin D, fatty acids, and protein) from 5,759 measurements of 731 fish species. Note that this dataset is expanded from the marine samples used in Hicks *et al. (Hicks et al. 2019)* (367 species) and includes data on freshwater species (Byrd et al. 2021) and new nutrient analyses of tropical coral reef species (Robinson et al. 2022a).

### Model structure

Fish consume nutrients according to their **diet (ecology), energetic demand (physiology/life-history)**, and **thermal regime (environment)**, in ways reflected by their individual traits (Budge et al. 2002, McGill et al. 2006, Mouillot et al. 2013). We used FishBase (Froese and Pauly 2010) to compile a set of 10 traits relating to these three key pathways (Figure 1).

**Figure 1.**
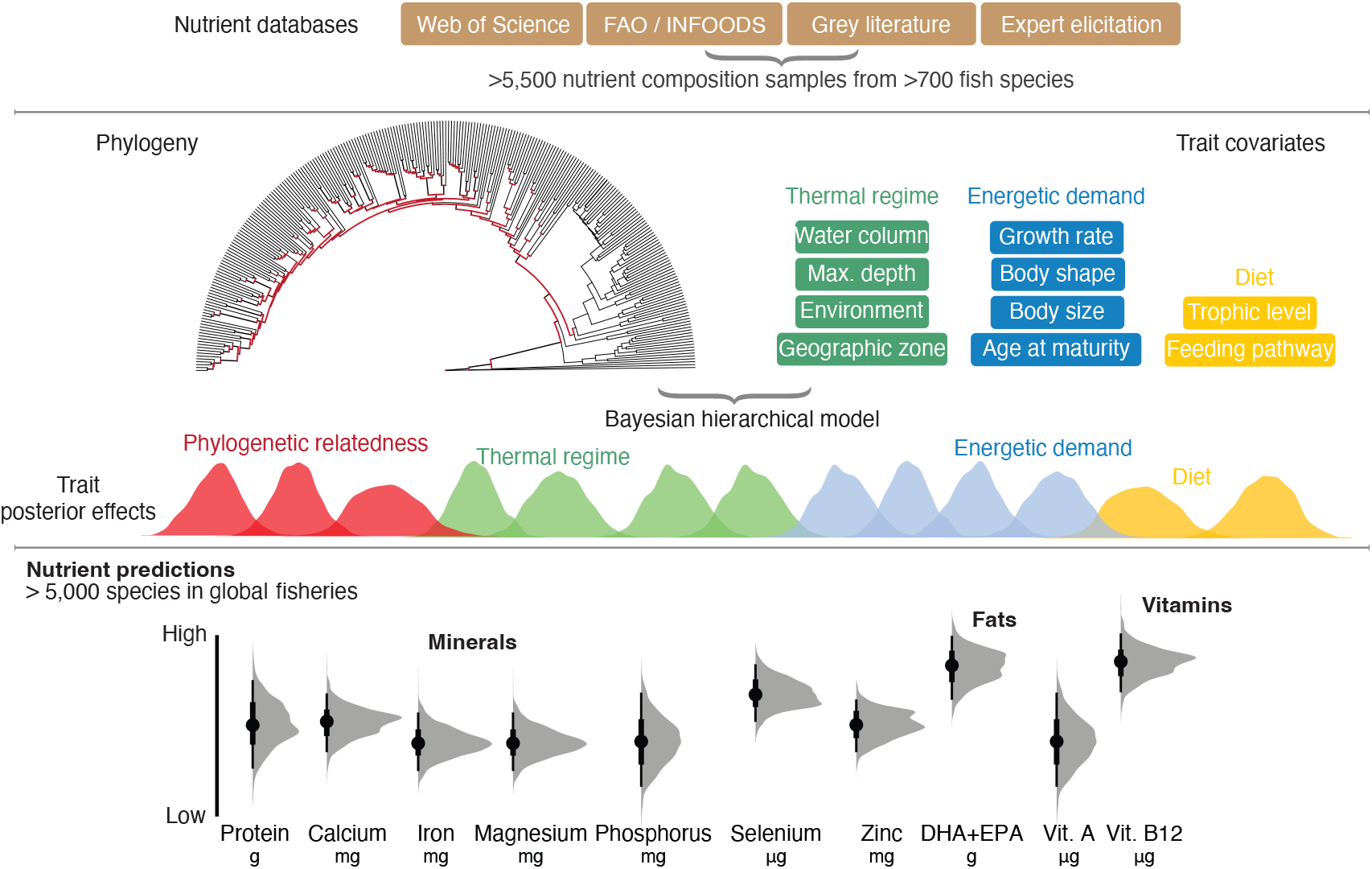
Conceptual figure of data sources, traits influencing nutrient content, model structures, and available outputs. For any given species, the lowest available phylogenetic unit is used (red distributions within the Bayesian hierarchical model). Grey distributions are full posterior densities for each nutrient in each species, with medians (black circles) and vertical lines representing 95% (thin lines) and 50% (thick lines) highest posterior density intervals.

**Diet** directly impacts the nutrient content of fish as these are the compounds that are ultimately synthesized into tissue (Willis and Sunda 1984, McGill et al. 2006). We represented diet through the traits of feeding pathway and trophic level. Feeding pathway (**FP**) refers to the main basal food sources in which dietary nutrients are sourced, through either the pelagic (*e*.*g*. planktonic feeding) or benthic (*e*.*g*. benthic, deep sea, demersal, algae) pathways (Table 1), that have distinct nutrient profiles based on distinct components of primary production and nutrient recycling (Pauly et al. 2000). Pathways for higher trophic level species reflected the feeding pathway of their prey or the dominant proportion of their prey’s feeding pathway for mixed-diet species. Trophic level (**TL**) refers to the number of feeding linkages between primary producers and a given species, which can reflect the bioaccumulation or bioreduction of specific nutrients (Van der Oost et al. 2003).

**Table 1.** Feeding pathway designation. Feeding pathways assigned (benthic or pelagic) based on FishBase functional group alternatives.

| Functional_group_I | Functional_group_II | Feeding_pathway |
| --- | --- | --- |
| detritus | Detritus | benthic |
| plants | Phytoplankton | benthic |
| plants | Other plants | benthic |
| zooplankton | Fish (early stages) | pelagic |
| zooplankton | Planktonic crustaceans | pelagic |
| zooplankton | Other planktonic invertebrates | pelagic |
| zoobenthos | Benthic crustaceans | benthic |
| zoobenthos | Cnidarians | benthic |
| zoobenthos | Echinoderms | benthic |
| zoobenthos | Insects | benthic |
| zoobenthos | Mollusks | benthic |
| zoobenthos | Sponges and tunicates | benthic |
| zoobenthos | Worms | benthic |
| zoobenthos | Other benthic invertebrates | benthic |
| nekton | Cephalopods | pelagic |
| nekton | Finfish | pelagic |

**Table 2.** Model comparison for out of sample predictive accuracy of fish nutrient trait (T) and trait-phylogeny (TP) models using leave-one-out cross-validation (LooCV) where Loo ELPD is the theoretical expected log pointwise predictive density for a new dataset.

| Nutrient | TP ELPD | TP SE | T ELPD | T SE | Diff (TP-T) |
| --- | --- | --- | --- | --- | --- |
| Vitamin D | -39.600000 | 8.100000 | -44.200000 | 6.000000 | 4.700000 |
| Vitamin B9 (Folate) | -4.900000 | 3.600000 | -12.200000 | 3.600000 | 7.300000 |
| Vitamin B12 | -60.100000 | 6.600000 | -72.600000 | 6.500000 | 12.500000 |
| Omega-3 | -318.800000 | 15.700000 | -347.000000 | 15.000000 | 28.200000 |
| Vitamin A | -433.700000 | 14.300000 | -481.200000 | 14.900000 | 47.500000 |
| Calcium | -984.800000 | 27.200000 | -1104.900000 | 44.400000 | 120.100000 |
| Iron | -938.100000 | 26.900000 | -1082.400000 | 28.000000 | 144.300000 |
| Selenium | -403.700000 | 26.400000 | -550.300000 | 28.800000 | 146.600000 |
| Magnesium | -157.400000 | 26.800000 | -337.600000 | 27.100000 | 180.100000 |

**Energetic demand** is a fundamental attribute of animals, reflecting trade-offs between growth and reproduction relative to somatic storage (Willmer et al. 2009). We represented energetic demand through the traits of maximum length (**L**_**max**_), age at maturity (**A**_**mat**_), the fish growth parameter (***K***), and body shape (**BS**). Maximum length scales directly with key attributes relating to home range size and metabolism, while age at maturity reflects the point at which resources are allocated to reproduction. The growth parameter *K* reflects the rate at which maximum body size is reached and therefore how energy is allocated to the accumulation of body mass. Body shape is indicative of how fish both feed and move through their environment, represented by being flat, elongate (or eel-like), fusiform, or having short-deep bodies.

**Thermal regimes** were identified to capture the metabolic demand (Brown et al. 2004) posed by living in colder environments as well geographical conditions such as terrestrial runoff that can impact nutrient availability (Barlow et al. 2018). Temperature covariates were maximum depth (**D**_**max**_) and other environmental conditions represented by four geographic zones (**GZ**): tropical, subtropical, temperate, and polar/deep. Environment (**EN**) refers to the aquatic regime, one of marine, freshwater, brackish, or mixed (*i*.*e*. more than one environment). Water column (**WC**) refers to the position in the water column where the species is typically found, one of pelagic, demersal, reef-associated, bathypelagic, or benthopelagic, each of which has distinct pathways for nutrient input and cycling.

While fish traits are directly linked to where and what fish eat, these characteristics are themselves correlated among closely related species, resulting in phylogenetically-predictable relationships for fish nutrient content (Vaitla et al. 2018). Therefore, we also included phylogenetic relatedness within the correlation structure of our statistical model, using a phylogenetic tree for bony fishes that included additional chondrichthyan relationships (see below).

Lastly, samples of fish tissue in our nutrients database included several sampling parameters – variables that influence sample collection but are not of direct interest – that we included to account for their potential bias relative to the nutrient content of edible fish muscle. These included the tissue type (muscle, whole, whole/parts, unknown; **FO**) and preparation (wet, dry, unknown; **PR**).

### Statistical models

To estimate the expected nutrient concentration of every species in our nutrient database, we developed two Bayesian hierarchical models for each nutrient: a modified version of the traits model used in Hicks *et al*. 2019 (‘traits’ model; T) and a new model with both trait and phylogenetic information present, the phylogenetic model having been used previously in predictive models of fish nutrients (Vaitla et al. 2018) and traits (Thorson 2020) (‘trait-phylogeny’ model; TP).

Previous analysis of fish nutrients by Vaitla *et al*. 2019 implemented phylogenetic least squares regression, which accounts for the phylogenetic dependence among species using covariance matrix weights, in their case using Pagel’s λ model to specify the weights (Vaitla et al. 2018). We adopted the related but more flexible 3/2 Matérn function (Williams and Rasmussen 2006), where the covariance between species *k* and *j* is given by a phylogenetic Gaussian Process:

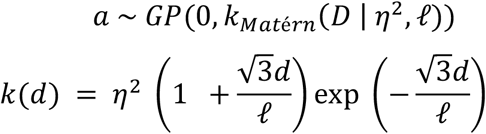

with priors

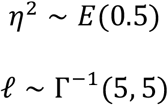

with parameters *η* and *ℓ* specifying an exponential rate of decline in correlation between species as the phylogenetic distance between them (*D*_*kj*_) increases. Our matrix of phylogenetic distances (*D*) was built by combining the time calibrated molecular phylogenies for ray-finned fishes (Actinopterygii; 31,526 tips) (Rabosky et al. 2018) with the phylogeny of cartilaginous fishes (Chondrichthyes; 1,198 tips; Stein et al. 2018). For each taxonomic phylogeny a single tree was randomly selected from the pseudo posterior tree distributions. The ray-finned fishes tree was added to the backbone chondrichthyan tree as a sister group at the root ∼425 MY using the phytools package in R (Revell 2024). Prior to analysis, trees were trimmed to the species of interest and scaled to a 0-1 correlation matrix of observed and predicted species. These relationships were then used as phylogenetic offsets relative to intercept *γ*_*_ in a linear model for each observation *i* that included a trait covariate (X)

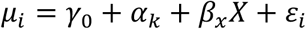

The set of species-level trait covariates were

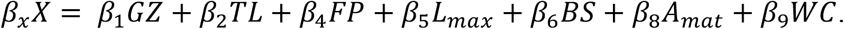

While we used a mix of Normal, Gamma, and Non-central *t* distributions for the data likelihood in Hicks *et al*. 2019, we chose to model nutrients in our trait-phylogeny (TP) model using a non-central *t* distribution

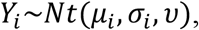

with an hetersocedastic error that accounted for differences in both sample preparation (wet, dry, unknown) and tissue type (muscle, whole, whole parts, unknown)

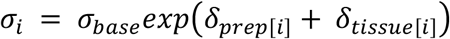

and weakly regularizing priors

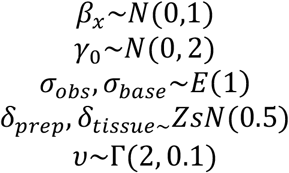

where *ZsN* is a zero-sum Normal distribution, where the sum of all categories is constrained to sum to zero.

We ran both models on each of the 12 nutrients, using the Python package PyMC (v5, www.pymc.io). Models were run with four separately initiated chains for 1,000 post-warmup iterations using a No-U-Turn sampler (NUTS). We examined posterior predictive distributions, expected cumulative distribution functions, and Gelman-Rubin statistics for evidence of model fitting problems for each nutrient and model combination (see https://github.com/mamacneil/NutrientFishBase). We also examined both the trait and trait-phylogeny models for model fit using leave-one-out cross validation (Loo-CV) and used these scores to compare them for out of sample predictive accuracy across nutrients (McElreath 2020).

### Predicting nutrients in fish globally

Our trait-phylogeny nutrient models provided a basis for predicting nutrients in any unmeasured species (*i*.*e*., ‘out-of-sample’ predictions). We extracted all fish species recorded in two global fisheries catch datasets (Sea Around Us Project and Illuminating Hidden Harvests) (Pauly 2007, Basurto et al. 2025) and two reef ecosystem monitoring programmes (Reef Life Survey and Social Ecological Research Frontiers) (Edgar and Stuart-Smith 2014, Cinner et al. 2020). These datasets represented 5,588 species, including the vast majority of reported marine and freshwater fisheries catches globally and most known non-cryptic fish species inhabiting coral reef ecosystems. We extracted traits for each species from FishBase, using family- and genus-level averages where species-level trait data were unavailable. We then predicted the nutrient content of all species for all 12 nutrients, reporting 50% and 90% highest posterior density intervals to reflect uncertainty in our estimates.

### Open-access nutrient predictions for global fishes

All species’ nutrient predictions were integrated with FishBase, an online, publicly available database of fishes. At https://www.fishbase.se/Nutrients/NutrientSearch.php, searching for specific species, genera, families, and geographies returns a table of the predicted nutrient concentration (with 95% uncertainty intervals) that can be downloaded as a csv, and raw values from the literature if they were available. Each species page also lists the predicted median nutrient concentration with uncertainty intervals. We further developed an application to communicate how model predictions can be linked to human health guidelines on recommended daily nutrient intakes. Our application (https://james-robinson.shinyapps.io/FishNutrientsApp/) visualizes the contribution of selected fish species to recommended daily intakes of calcium, iron, selenium, zinc, total omega-3 fatty acids, and vitamin A. We estimated nutrient contributions for ‘wet’ fillet samples and several dietary populations (children 6 months - 5 years old, adult women and men, and pregnant women), that scale with portion size. Nutrient reference values were extracted from harmonized guidelines (Allen et al. 2020, Robinson et al. 2025), and all nutrient contributions were capped at 100%. The application was written using Shiny in R (Chang et al. 2025).

## Technical Validation

Model convergence was assessed using the Gelman-Rubin diagnostic (R-hat < 1.01 for all parameters) and effective sample size (ESS > 400 for all parameters). Posterior predictive checks (Figure 3) confirmed that the TP model adequately captured the observed data distribution for all nutrients. Model calibration was assessed by the proportion of observations falling outside 95% highest posterior density intervals (Figure 2); a well-calibrated model should have approximately 5% of observations outside these intervals. Leave-one-out cross-validation using Pareto-smoothed importance sampling (PSIS-LOO) demonstrated superior out-of-sample predictive performance for the TP model compared to the T model for all 12 nutrients (LOO ELPD differences ranged from 4.7 for Vitamin D to 459 for Phosphorus)..

**Figure 2.**
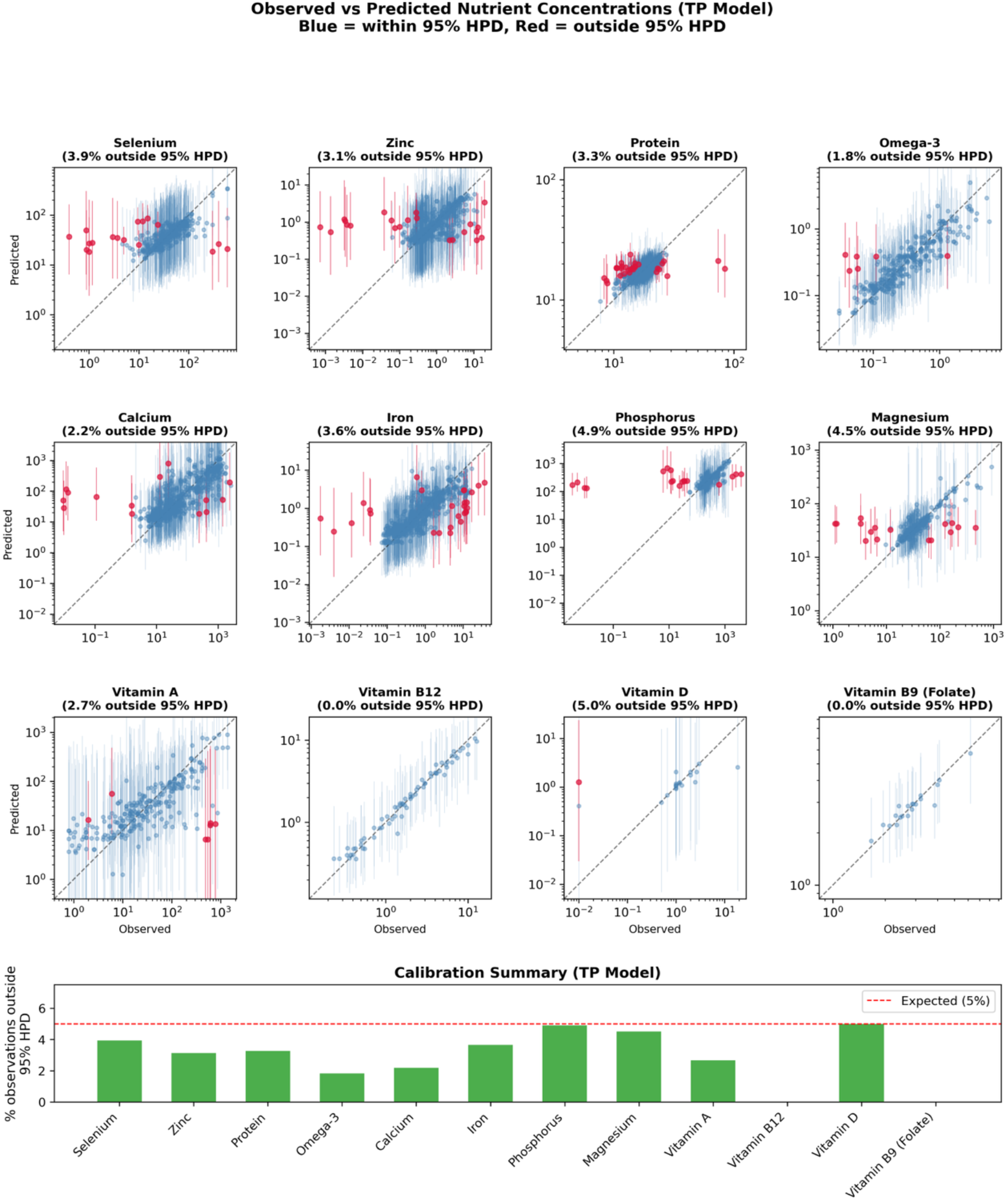
Posterior predicted data-distribution overlaid with observed datapoint for each nutrient under the trait-phylogeny (TP) model. Observations falling outside their corresponding 95% highest posterior density interval are plotted in red. Bottom panel histograms of percentage observation outside of 95% HPD per nutrient.

**Figure 3.**
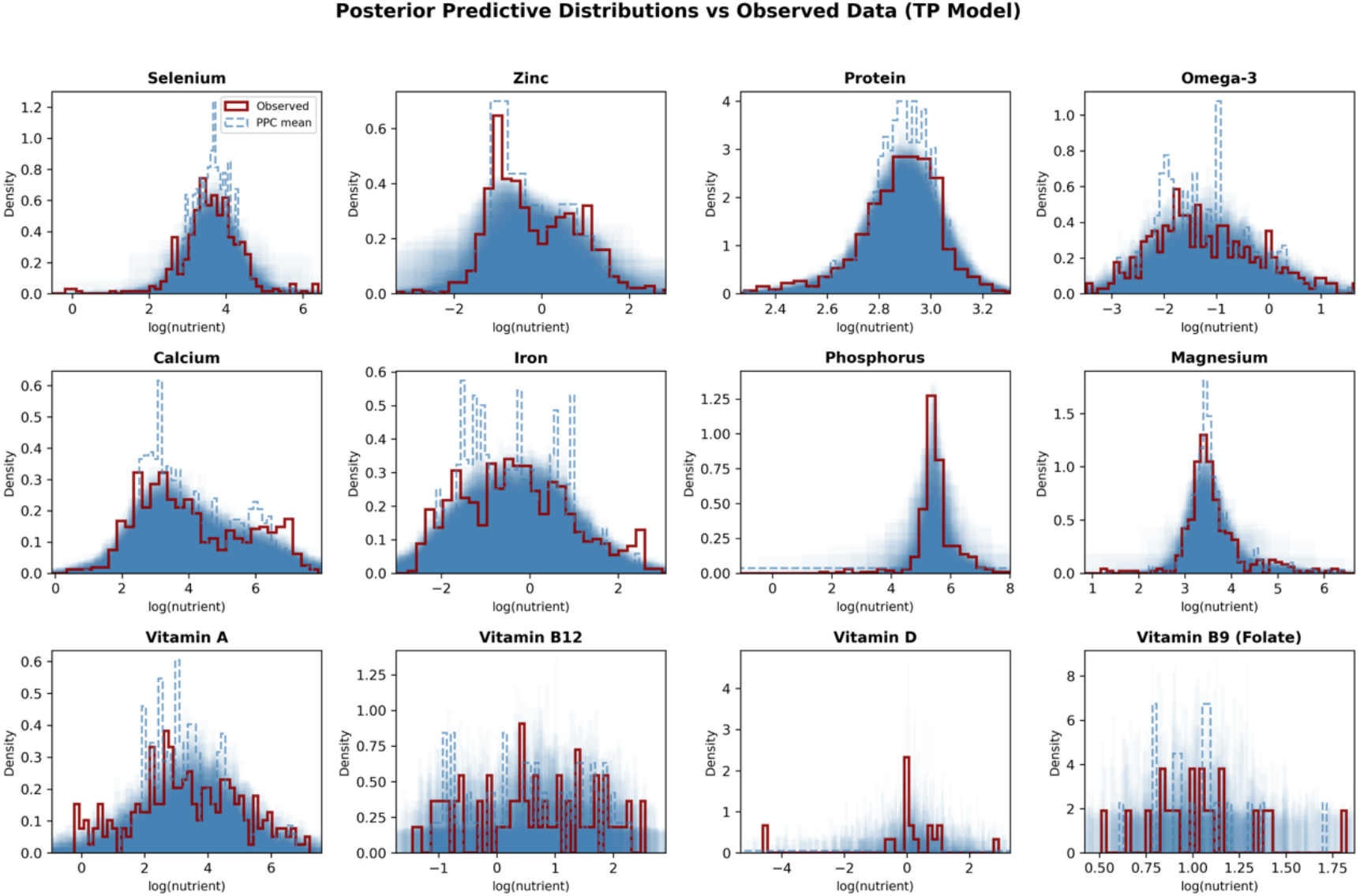
Posterior realizations (blue) of observed data (red) across nutrients under the trait-phylogeny (TP) model.

**Figure X.**
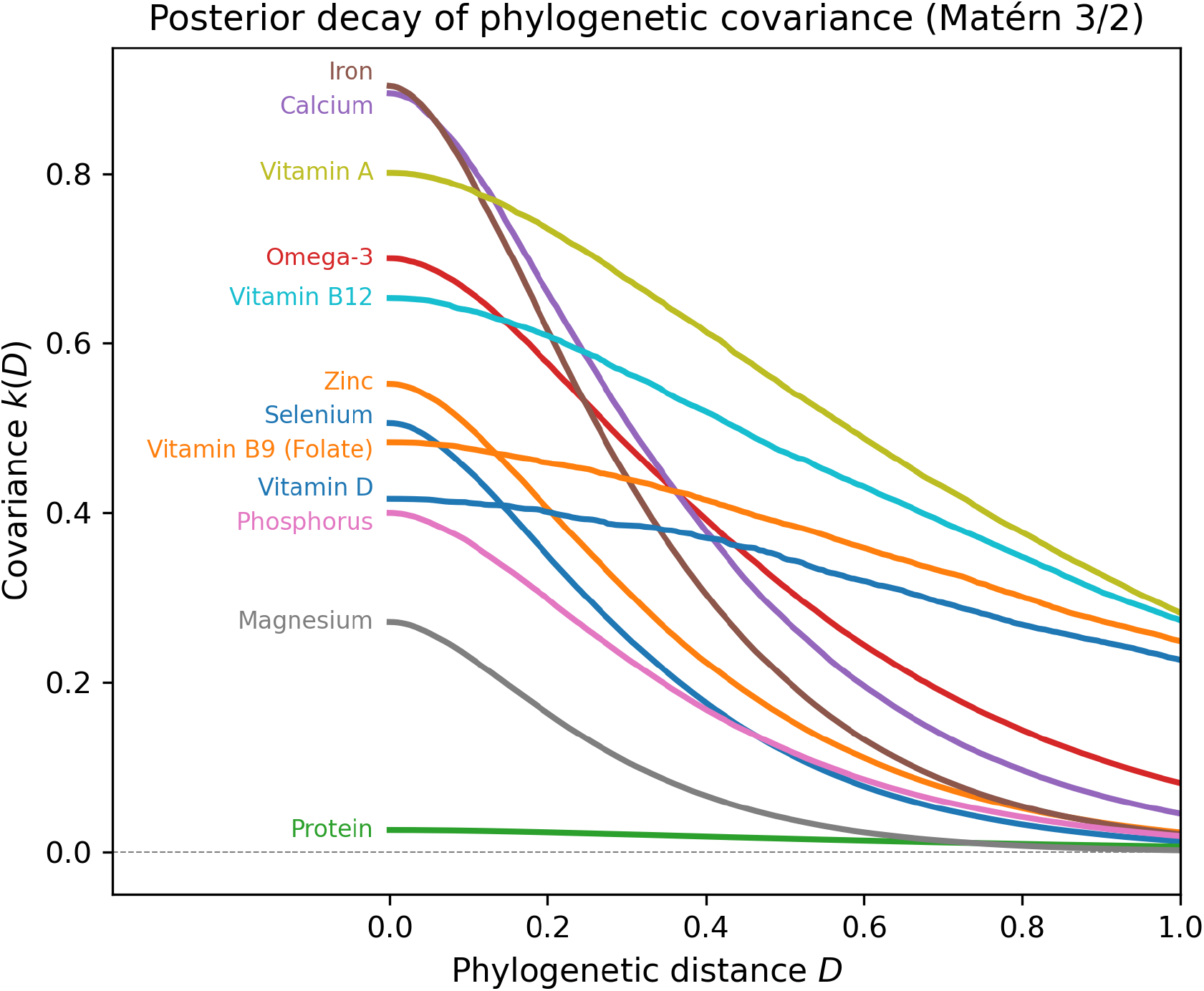
Phylogenetic decay in nutrient concentration covariance among fishes given a Matérn 3/2 kernel density function for the trait-phylogeny (TP) model. Phylogenetic distances are scaled relative to maximum distance in the data (1).

## Data Availability

All data records associated with this work are available through a GitHub repository (https://github.com/mamacneil/NutrientFishBase). The observed nutrient concentration database (D1: all_nutrients_active_2025.xlsx) contains 5,759 measurements from 731 species across 12 nutrients. The pairwise phylogenetic distance matrix (D2: fish_shark_phylo_distances.csv) is available via Dropbox (https://www.dropbox.com/scl/fi/ye2rterorz0rwv199a8cc/fish_shark_phylo_distances.csv?rlkey=vi8wsxal9pk3zk4xhu47r94uq&dl=0). MCMC posterior samples for the trait-phylogeny (M1) and trait (M2) models are stored as NetCDF-4 files in model/Traces/. Model comparison results (M3: model_comparison_all_nutrients.csv) and species-level nutrient predictions for 5,588 species across 12 nutrients (P1: fishbase_predictions_2026.csv) are available in the data/ directory. Predicted nutrient concentrations are also publicly accessible through FishBase (https://www.fishbase.se/Nutrients/NutrientSearch.php).

## Code Availability

The code used to generate these data records is available at https://github.com/mamacneil/FishNutrients. The analysis pipeline is implemented in two Jupyter notebooks: fit_all_nutrients_full_pipeline.ipynb (model fitting, comparison, and per-nutrient diagnostics) and generate_figures.ipynb (summary figures and tables). The analysis was performed using Python 3.11.6 with PyMC 5.10 for Bayesian inference and ArviZ 0.16.1 for posterior analysis. Claude Code Opus 4.6 was used to annotate and streamline code for production upload to GitHub.

## Author contributions

M.A.M. Designed the study, developed and implemented modelling, and wrote the initial draft manuscript.

E.M. Collected data, developed modelling and contributed to the manuscript.

J.P.W.R. Collected data, developed modelling and contributed to the manuscript.

N.A.J.G. Contributed to the model conception and manuscript.

P.J.C. Secured funding, designed the study, and contributed to the manuscript.

M.L.D.P. designed the study and contributed to the manuscript.

C.C.H. Secured funding, designed the study, and contributed to the manuscript.

## Acknowledgements

Thanks to A Andorra for public PyMC code used in our initial models and C Mull for help merging phylogenies. We acknowledge M. Roscher, C. D’Lima, N. Swan, J. Silveira, S. Masangwi, A. P. Belsan, S. Silver, D. Raz, B. Wisman and S. Das for their contributions to the compilation of the original databases, and S. Thilsted for support and advice.

